# Identification of HIV-1 infected cells in single-cell multiome datasets using Kraken2

**DOI:** 10.64898/2026.11.27.767303

**Authors:** Lidia Garrido-Sanz, Victor Urrea, Ramon Lorenzo-Redondo, Leah B. Soriaga, Maria C. Puertas, Judith Dalmau, Pep Coll, Beatriz Mothe, Christian Brander, Bonaventura Clotet, Amalio Telenti, Javier Martinez-Picado, Sara Moron-Lopez

**Affiliations:** IrsiCaixa, Badalona, Barcelona, 08916, Spain; Feinberg School of Medicine, Northwestern University, Chicago, Illinois, 60611, United States; Trail Biomed, Lausanne, Switzerland; Centro de Investigación Biomédica en Red de Enfermedades Infecciosas (CIBERINFEC), Instituto de Salud Carlos III (ISCIII), Madrid, 28029, Spain; Germans Trias i Pujol Research Institute (IGTP), Badalona, 08916, Spain; BCN CheckPoint - Projecte dels NOMS-HISPANOSIDA, Barcelona, 08015, Spain; Department of Infectious Diseases and Fundació Lluita contra les Infeccions, Hospital Universitari Germans Trias i Pujol, Badalona, Barcelona 08916, Spain; Catalan Institution for Research and Advanced Studies (ICREA), Barcelona, 08010, Spain; Department of Infectious Disease and Immunity, University of Vic-Central University of Catalonia (UVic-UCC), Vic, 08500, Spain

**Keywords:** ATAC-seq, RNA-seq, bioinformatics, HIV-infected cells, Cell Ranger, virKra

## Abstract

1.

Detecting HIV-1-infected cells in single-cell sequencing studies is challenging primarily due to low transcriptional activity, scarcity of infected cells, and high viral genetic variability. Here, we present a novel pipeline integrating multiple HIV-1 subtype B genomes with Kraken2 (K2) for k-mer-based classification, bypassing the need for exact sequence alignment. Peripheral blood mononuclear cells from five individuals were analyzed at three time points: before HIV acquisition, at the time of HIV diagnosis, and after one year on antiretroviral therapy (ART). Single-cell multiome libraries - comprising nuclei ATAC-seq (ATAC) and gene expression (GEX) data-were processed using K2 against a curated HIV-1 subtype B reference library. Calibration was performed using pre-HIV acquisition samples, and optimized parameters applied to post-HIV and post-ART samples. For comparison, the standard Cell Ranger-ARC (CR) pipeline was also used. Out of 150,060 single nuclei analyzed, our pipeline identified 32 HIV-infected cells (13 in ATAC datasets, 18 in GEX datasets, and 1 in both), whereas CR detected 11 infected cells exclusively in GEX and yielded one false positive. This innovative bioinformatic approach enhances the detection of HIV-infected cells compared to CR and may complement studies aimed at defining cellular markers identifying infected cells.

**Highlights:**

- A k-mer-based pipeline detects HIV-1-infected cells in multiomics single-nuclei data.
- 1,312 HIV-1 subtype B genomes were used to populate the viral reference database.
- Pre-infection samples were used to curate viral references and calibrate the pipeline.
- It found 32 infected cells in 150,060 nuclei, versus 11 (10 + 1 FP) with the standard pipeline.

**Graphical abstract:** 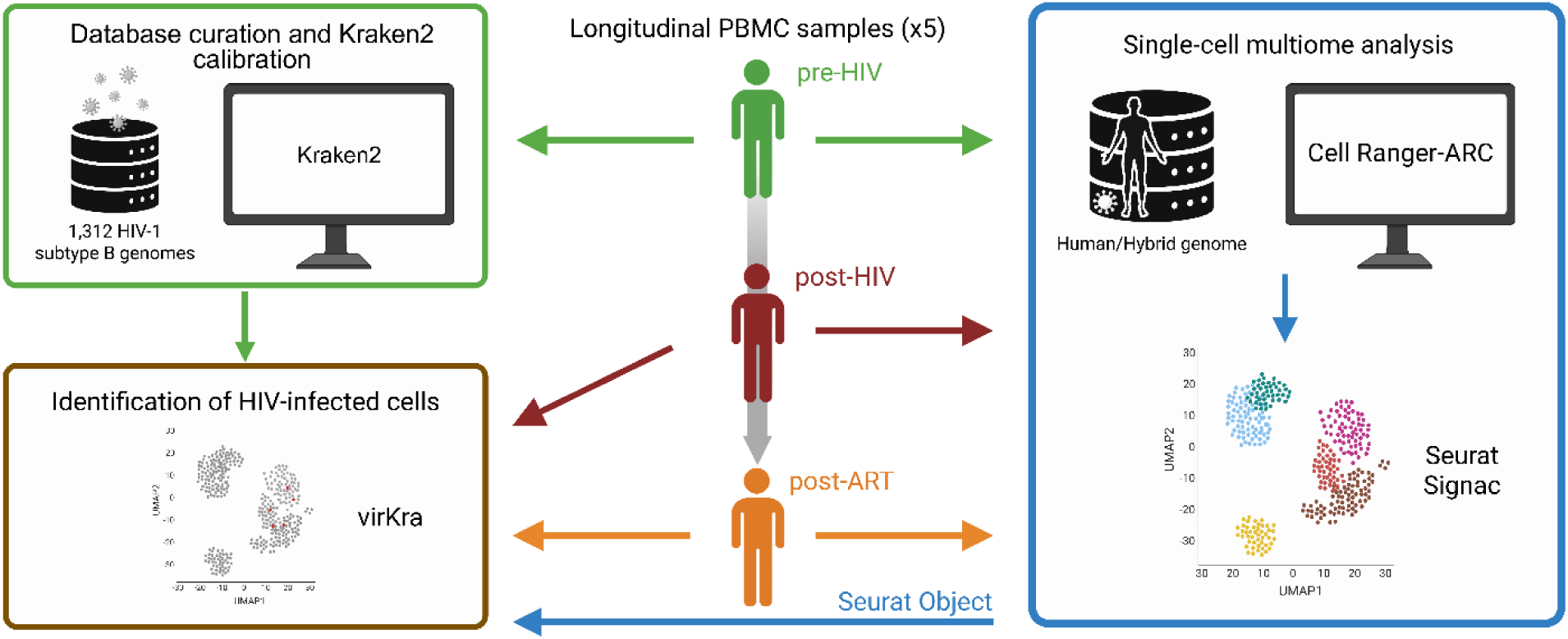

## 2. Background

Human immunodeficiency virus type 1 (HIV-1) continues to pose a significant global health challenge, despite more than four decades of intensive research and the widespread use of antiretroviral therapy (ART) (1). HIV-1 displays remarkable genetic diversity, primarily driven by the high mutation rate of its reverse transcriptase enzyme and rapid replication dynamics (2, 3). This extraordinary genetic variability is classified into four major groups: M, N, O, and P (4, 5); among these, group M is the primary lineage responsible for the global HIV epidemic. Group M is further subdivided into nine distinct subtypes and numerous circulating recombinant forms, each with characteristic geographic distributions, being subtype B the most predominant in the Americas, Europe, and Oceania (6). These subtypes typically exhibit inter-subtype genetic distances of approximately 10-20% (7), reflecting the evolutionary history of the virus as well as distinct epidemic patterns across global regions. This diversity has important implications for diagnosis, treatment, and vaccine development.

Recent advances in single-cell transcriptomic and epigenomic technologies, combined with state-of-the-art bioinformatics, have significantly propelled HIV-1 research by enabling the detailed characterization of both infected and bystander cells at an unprecedented resolution (8, 9). However, accurately identifying HIV-1 DNA from integration sites within the host genome, as well as HIV-1 RNA sequences arising from active transcription at the single-cell level, remains challenging. These difficulties stem from the extraordinary genetic variability of the virus, the limited abundance of infected cells (even in samples enriched for CD4^+^ T cells; 10, 11) and low transcriptional activity, particularly in people with HIV (PWH) under ART virological suppression (12–16). This challenge is critical because most HIV-1-detection tools currently used in single-cell studies are devised for eukaryotic genomes and depend on exact sequence matches when comparing query reads to references (17, 18), a strategy that may fail due to the vast viral genetic diversity. To improve detection efficiency with these tools, three general strategies are typically applied, either independently or in combination: (i) using autologous HIV-1 reference sequences (a labor-intensive, complex, and costly approach that nevertheless ensures precise representation of individual-specific viral variants; 19); (ii) incorporating multiple diverse viral genomes within the reference, which circumvents the need for individualized references by expanding the genetic variability represented and requires careful curation to avoid incorrect sequences (16, 20); and (iii) relaxing aligner parameters to increase the number of classified reads at the expenses of potentially increasing false positives (21). Alternatively, specialized classification tools designed for viral detection have been developed to enhance sensitivity in identifying viral sequences (22–26).

Because currently available pipelines and tools for single-cell experiments aimed at identifying HIV-1 DNA or RNA are suboptimal, we present here a robust and generalizable bioinformatic pipeline designed to accurately detect HIV-1-infected cells in single-cell datasets. Our approach incorporates multiple HIV-1 subtype B viral genomes to create a curated reference database and employs a k-mer-based tool, Kraken2 (K2) (27) for the detection of infected cells, capable of identifying both viral DNA integrated in the host genome and viral RNA from various stages of the viral replication cycle. K2 is a rapid, memory-efficient software that classifies sequenced reads by matching short genomic fragments (k-mers) to a database, enabling taxonomic resolution of target organisms. It uses a compact hash table and stores “minimizers” of k-mers alongside their associated taxon for precise and efficient classification of metagenomic data using the Lowest-Common Ancestor algorithm. For enhanced accuracy, both the reference database and K2 are calibrated iteratively using samples collected from five individuals prior to HIV acquisition. The optimized K2 and curated references are subsequently applied to longitudinal samples from the same individuals after HIV acquisition before and one year after initiation of ART. For benchmarking, all samples were also processed using Cell Ranger-ARC (28, 29) the standard pipeline for single-cell multiome data analysis.

## 3. Materials and Methods

### 3.1. Samples datasets

For this analysis, samples described in HIV-scAR project (30) were utilized. In that study, longitudinal peripheral blood mononuclear cell (PBMC) samples were collected from five individuals at three distinct time points: prior to HIV-1 acquisition (pre-HIV), at the time of HIV-1 diagnosis (post-HIV), and after one year on ART (post-ART). These samples are particularly valuable because PBMCs from each time point were obtained from the same individuals. The inclusion of pre-HIV samples enables optimization of software parameters and refinement of reference datasets, thereby enhancing the accuracy of read-to-reference assigning.

The detailed library preparation and sample processing are described in HIV-scAR (30). Briefly, nuclei were isolated from 1,000,000 PBMCs for each sample following the 10x Genomics protocol (CG000365 Rev. C). Libraries for single-nucleus assay for transposase-accessible chromatin (ATAC) and gene expression (GEX) were generated using “Chromium Next GEM Single Cell Multiome ATAC + Gene Expression” (CG000338 Rev. E). Sequencing was performed on an Illumina NovaSeq 6000 system, generating paired-end reads of 49-50bp for ATAC and single-end reads of 90 bp for GEX. FASTQ files underwent quality control analysis with FastQC (v0.11.9) (31). High-quality FASTQ files were aligned to the human genome GRCh38.p12 (from the 10x Genomics support repository: refdata-cellranger-arc-GRCh38-2020-A-2.0.0.tar.gz) using Cell Ranger-ARC (cellranger-arc-2.0.2, http://support.10xgenomics.com/) with the “count” option. Resulting peak-barcode and gene-barcode matrices were filtered with Seurat (v4.3.0) (32) and Signac (v1.9.0) (33) in R (version 4.3.0) (34). The features matrices were cleaned from background contamination and empty droplets with SoupX package (v1.6.2) (35), and doublets were detected and removed with the scDblFinder package (v1.14.0) (36). High-quality cells were retained by applying 1.5 Interquartile Range (IQR) thresholds to the number of genes, UMI counts, and ATAC molecules. We also removed cells with transcription start site (TSS) enrichment below 1, nucleosome signal above 2, or mitochondrial percentage above 20%. Remaining cells were loaded into a Seurat Object. GEX data was normalized using the SCTransform() function, and individual samples were integrated with reciprocal principal component analysis (RPCA). ATAC data underwent peak calling with MACS2 (37) and normalization with RunTFIDF(), which applies the term frequency-inverse document-frequency (TF-IDF) method (38). Both GEX and ATAC data were integrated using the Weighted Nearest Neighbor (WNN) graph. The integrated multimodal data were clustered at a resolution of 0.8, annotated with Azimuth level 1 reference (32), and plotted using Uniform Manifold Approximation and Projection (UMAP) for non-linear dimensional reduction. Further details of HIV-scAR project are available in (30).

### 3.2. HIV-1 reference genomes

In December 2022, we retrieved all complete HIV-1 subtype B genomes from the European geographic region available in the LANL HIV Database (https://www.hiv.lanl.gov/). Based on these criteria, a total of 1,312 complete genomes were obtained (Supplementary Table S1).

To facilitate the identification of viral sequences, we constructed a hybrid human-viral genome by concatenating the human genome FASTA and GTF files with the viral genome files. Since our primary objective was to detect viral sequences rather than pinpoint the exact location within the HIV-1 genome, we generated custom annotations by designating entire viral genomes as individual genes. This approach enables computational tools to recognize any viral read, regardless of its genomic position, thereby increasing the number of viral sequences detected. Aligners-based tools are designed to assign reads unambiguously to a single reference genome. Based on this principle, we also evaluated whether expanding the reference database to include multiple closely related viral genomes, representing viral quasi-species, would impact the number of HIV sequences identified. For comparison, the human genome was concatenated with the HXB2 viral reference genome (accession number K03455).

### 3.3. Bioinformatic pipeline

In this study, we selected two bioinformatic tools designed to associate sequencing reads with reference genomes, each employing distinct methodological paradigms: (i) an alignment-based approach, which aligns entire reads to a reference sequence, and (ii) a k-mer-based approach, which assigns reads to specific taxa based on short sequence fragments. For the alignment-based method, we chose Cell Ranger-ARC (CR), a gold-standard pipeline for multiome analysis that uses BWA-MEM for ATAC data alignment (17) and STAR software for GEX data (18). For the k-mer-based approach, we selected Kraken2 (K2) (27), as this tool has been extensively evaluated and benchmarked against other genomic classification methods (39, 40).

Despite the high complexity of the underlying algorithms implemented in the aforementioned software, none of them (when executed with default parameters) ensures optimal performance (41). In other words, no tool provides results entirely free from false positive detections, such as reads wrongly assigned to a reference genome. To improve result accuracy, K2 was calibrated using pre-HIV samples as negative control, since these samples are devoid of HIV-1 virus (Fig. 1). In practice, only K2 was calibrated, as its optimization involves adjusting a single parameter (the confidence score) which is both rapid and straightforward. In contrast, BWA and STAR possess multiple tunable parameters, making their calibration time-consuming and computationally demanding. Furthermore, parameter accessibility is limited, as BWA and STAR options cannot be directly modified through CR command line arguments.

**Figure 1.**
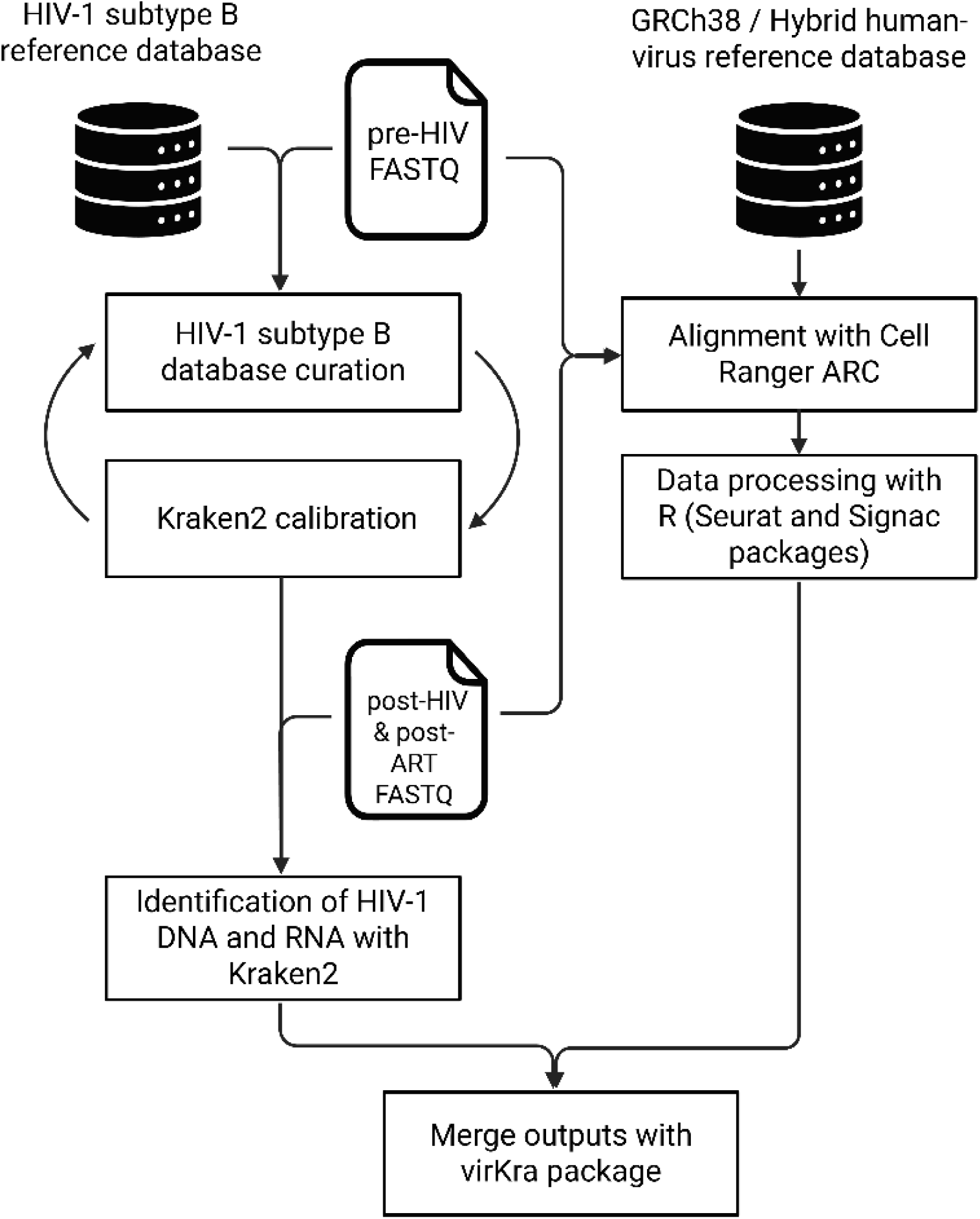
Bioinformatic pipeline run in this analysis. This figure has been generated with BioRender.

Reference databases that have not been thoroughly curated can impede the calibration process, as contamination and assembly errors may introduce extraneous sequences into reference genomes, leading to false positive (FP) detections (16, 20, 42, 43). In line with the calibration approach, non-curated genomes present in our viral reference database can be efficiently identified using pre-HIV samples and subsequently excluded to preserve the integrity of the database.

Calibration of the software and refinement of the reference database were conducted with K2 using an iterative strategy to ensure the robustness of tool parameters and the reliability of the reference genomes. To optimize read-to-reference assignments, we implemented a two-phase approach. In the first phase, we defined a parameter search space by specifying a range of values for each adjustable parameter of the classification software, aiming to identify configurations that yielded zero FPs. If all parameter sets produced FPs, a second phase was initiated to identify problematic references (defined as HIV-1 sequences that matched query reads from pre-HIV samples). Reads resulting in FP classifications were recovered and aligned using NCBI BLASTn (https://blast.ncbi.nlm.nih.gov/Blast.cgi, accessed in April 2023) (44, 45) against the “Nucleotide collection (nr/nt)” database, targeting HIV-1 organisms. From the resulting alignments, accession numbers corresponding to the matching references were extracted, and any references present in our viral reference database were removed. The refined database was subsequently used in a new round of read classification employing the same parameter combinations (Fig. 1). To further characterize the problematic references, we also aligned them with NCBI BLASTn against the “Nucleotide collection (nr/nt)” database.

#### 3.3.1. Taxonomic classification with Kraken2

Kraken2 (v2.1.0) was executed with the following options: --db, --confidence, --use-names, --gzip-compressed, --threads (set to 16), --classified-out, and --output for GEX dataset; additionally, the -- paired option was applied for the ATAC dataset. Calibration was performed by testing a range of confidence scores (using the --confidence option), from 0 to 0.50 in increments of 0.05, separately for ATAC and GEX datasets. Notably, K2 classifies paired-end reads as a pair, meaning it uses the k-mer information from both reads to assign either both reads of the pair to a taxon or neither. Next, post-HIV and post-ART samples were analyzed using K2 with optimized confidence scores. Reads classified as viral were linked to cells using 10x barcodes. For this step, we developed an in-house R package, virKra (v1.0.0; available on GitHub: https://github.com/irsi-grec/virKra), which extracts the names of reads mapping to the virus and retrieves their barcodes (from the i5 index read for ATAC and from R1 for GEX), as well as the UMI sequence for GEX. The barcode and sample information are then used to import this data into the Seurat object; for GEX, UMI sequences are additionally used to deduplicate reads replicated by PCR.

In single-cell multiome datasets, high-quality cells are selected through comprehensive bioinformatic pre-processing. This workflow typically involves the removal of low-quality cells, dead cells, empty droplets, and doublets (16). Rigorous cell filtering effectively eliminates most poor-quality cells, therefore, we reasoned that calibration of K2 could be performed by examining the DNA/RNA viral counts in the Seurat matrices (referred here as the CK2-SO) as an alternative to the previous calibration approach, which we denote as the CK2-FQ. We anticipate that tuning K2 using this strategy may allow for a lower confidence score threshold to remove FPs while increasing the number of robust viral DNA and RNA identifications. To implement the CK2-SO calibration, we defined a smaller search space for the confidence score, ranging from 0 up to the confidence score fixed by the CK2-FQ calibration (since K2 does not report assignments beyond those thresholds).

#### 3.3.2. Alignment with Cell Ranger-ARC

All multiome FASTQ files were aligned to the hybrid human-viral reference genome using the CR’s count option (cellranger-arc-2.0.2, http://support.10xgenomics.com/). The resulting output matrices, containing both transcript and peak count information, were processed with the same bioinformatic pipeline and software described previously for the HIV-scAR project. The entire workflow was also repeated using a hybrid reference consisting of the human and the HXB2 viral genome.

### 3.4. Evaluation metrics

The performance metrics used to evaluate the computational tools included the number of FP HIV-positive reads and cells, concordance correlation of infected cells, and execution time. In this study, we define these metrics as follows and compute them using specific data types according to their intended purpose:

- False positives (FPs): The number of reads incorrectly assigned to the virus and cells misidentified as infected in pre-HIV samples.
- Concordance between expected and observed proviral counts: Assessed using Bland-Altman analysis. We compared the total HIV DNA copies per million cells to the total count of HIV DNA/RNA (calculated as the sum of viral ATAC paired-end reads and GEX UMI counts) per million cells. Only post-HIV and post-ART samples were included in this analysis.
- Execution time: The total time required to run the K2 and virKra algorithms. The execution time for CR was not considered, as it is necessary for generating the Seurat Object.

## 4. Results

### 4.1. Viral reference curation and Kraken2 calibration

Because reference genomes may contain extraneous sequences arising from misassembly or contamination, which can lead to false-positive (FP) detections, we first assessed the integrity of the viral reference database using pre-HIV samples as negative controls during calibration of Kraken2’s (K2) confidence score. Calibration with all 1,312 HIV-1 subtype B genomes from the LANL HIV Database repository yielded FPs in all pre-HIV samples across all tested confidence scores both in ATAC (Fig. 2A) and GEX datasets (Fig. 2B). Subsequent analysis with NCBI BLASTn identified six conflictive genomes (Supplementary Table S1). The length of the non-viral strings ranged from 30 to 387 nucleotides and were located at either the 5’ or 3’ ends of the genomes. Misassembled genomes were removed from the database, and the remaining genomes were deduplicated using Seqkit rmdup (v2.5.1) (46), resulting in a total of 1,163 viral genomes for further analysis.

**Figure 2.**
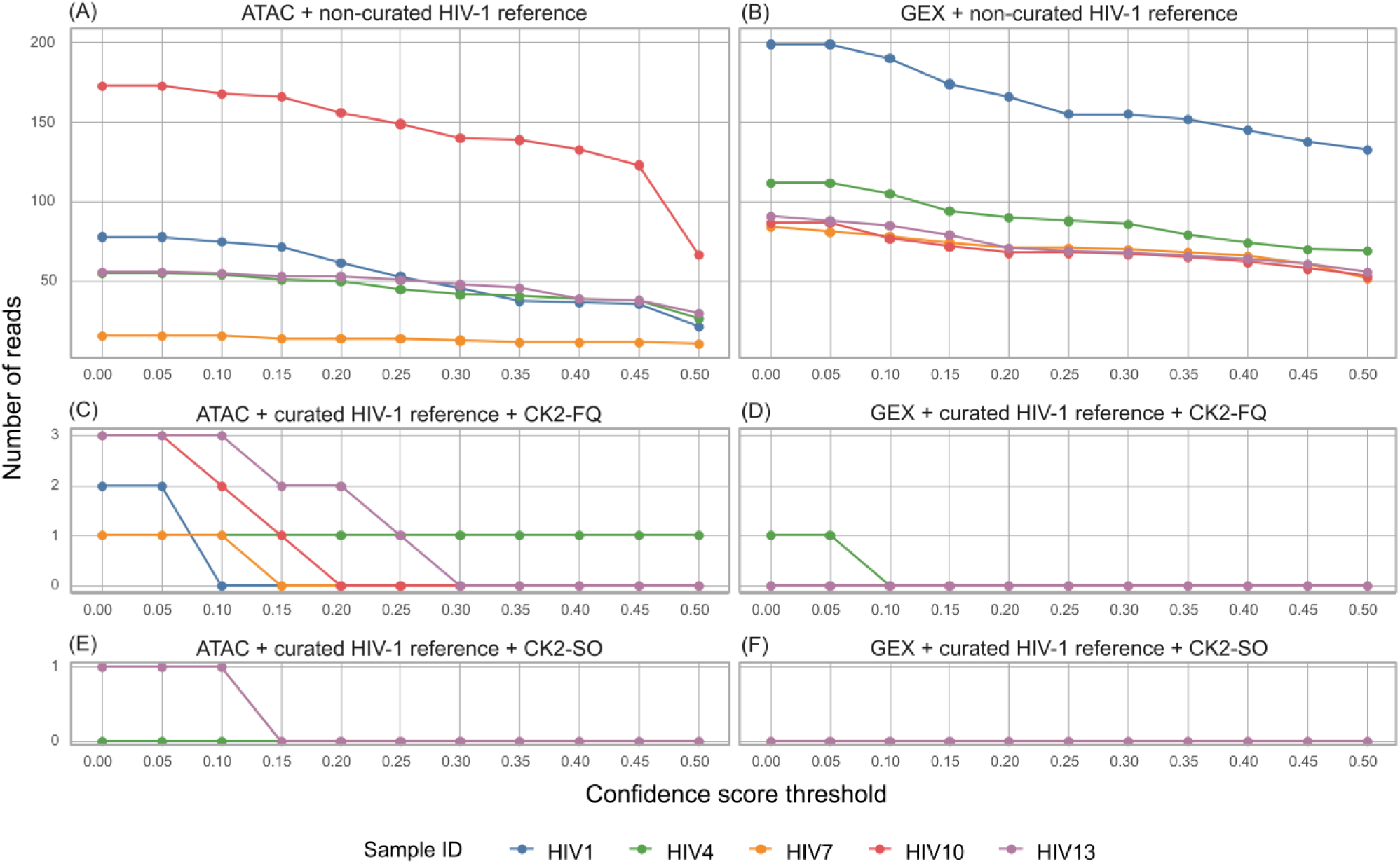
Number of reads assigned to HIV-1 in pre-HIV samples during K2 calibration and reference refinement. Colors indicate samples ID. Initially, pre-HIV samples were analyzed using K2 with a range of confidence scores (0 to 0.50, in increments of 0.05) applied to a non-curated viral reference database. After identifying and excluding problematic reference genomes, FASTQ-level calibration (CK2-FQ) was performed based on K2’s default output, followed by Seurat Object-level calibration using results transferred into the Seurat Object (CK2-SO). In this figure, the top row displays CK2-FQ calibration results for K2 using the non-curated reference database for ATAC (A) and GEX (B) datasets. The middle row presents CK2-FQ calibration results for ATAC (C) and GEX (D) datasets after reference refinement. The bottom row shows CK2-SO calibration results for ATAC (E) and GEX (F) datasets. For ATAC data, the number of reads refers to the count of read pairs classified as viral, while for GEX data, it refers to the count of single-end reads classified as viral.

Using the curated reference database, we calibrated K2 for multimodal datasets. In ATAC samples, applying a confidence score threshold of 0.30 eliminated all method-related FPs (Fig. 2C); the only remaining read pair (HIV4) was genuinely viral and attributable to index hopping (see Discussion). Although the reference curation step did not remove this read pair, subsequent NCBI BLASTn analysis confirmed its definitive alignment to HIV-1 genomes. In GEX samples, reference genomes cleaning successfully removed false positives across all samples at a confidence score of 0.10 (Fig. 2D).

Recognizing that pre-processing steps may remove cells failing quality control, we proceeded to calibrate K2 by examining the classified sequences retained within the Seurat Object. Using this CK2-SO strategy, we identified three pairs of reads from the ATAC dataset, which were excluded by applying a confidence score threshold of 0.15; no reads were detected in the GEX dataset (Fig. 2E-F and Supplementary Table S2). As a result, the CK2-SO approach enabled us to lower the optimized confidence score thresholds to 0.15 for ATAC and 0 for GEX.

In summary, our findings demonstrate that the use of default software parameters and non-curated viral genome references leads to FP detections of viral DNA and RNA. However, we show that reference curation and parameter optimization, implemented through our CK2-FQ strategy, effectively eliminate all FPs. Furthermore, the CK2-SO approach leverages cell-level quality control performed with the Seurat and Signac R packages, enabling more relaxed confidence score thresholds while maintaining accuracy.

### 4.2. Evaluation of HIV-1-positive cells identified by Kraken2

To evaluate the capacity of each pipeline to identify HIV-1 positive cells, we subsequently ran K2 with parameters optimized through the CK2-FQ calibration strategy on post-HIV and post-ART samples. HIV positive cells were detected in all five post-HIV samples, but only in three post-ART samples (Fig. 3A-B). Viral RNA was consistently present across these samples, whereas viral DNA was identified in only five samples (four post-HIV and one post-ART). The number of infected cells varied substantially between samples, with higher counts observed in post-HIV samples. Specifically, post-HIV samples contained an average of 2.2 ± 1.5 (format: mean ± standard deviation) HIV-positive cells per sample identified via ATAC-seq, and 8.8 ± 9.7 via GEX profiling (out of 9,573.6 ± 2,888.2 cells per sample). In contrast, post-ART samples showed lower numbers, with 0.4 ± 0.9 HIV-positive cells detected by ATAC-seq and 0.8 ± 0.8 by GEX (out of 9,937.6 ± 1,499.2 cells per sample). Viral DNA and RNA counts also differed across samples (Fig. 3C-D) remaining higher in post-HIV samples, where we detected 3.6 ± 2.9 DNA counts and 33.6 ± 64.5 RNA counts per sample (with 1.6 ± 1.0 DNA counts and 3.8 ± 17.8 RNA counts per cell). In post-ART samples, 0.6 ± 1.3 DNA counts and 0.8 ± 0.8 RNA counts were observed (with 1.5 ± 0.7 DNA counts and 1.0 ± 0.0 RNA counts per cell). Interestingly, sample HIV8 (post-HIV from participant 3, collected within the first 25-30 days after HIV acquisition) contained 149 RNA counts from 26 cells, with most counts arising from a single cell with 119 RNA molecules (Supplementary Tables S2 and S3). To evaluate concordance between proviral and estimated viral counts, we aggregated the viral DNA and RNA counts (Supplementary Table S4). The analysis revealed a discrepancy for sample HIV8, while the remaining samples clustered around the mean, indicating that our method produces concordant results (Supplementary Fig. S1A-B). After transferring CK2-FQ results into the Seurat Object from HIV-scAR project (named as CK2-FQ+SO) (Fig. 4B), 33 out of 60 HIV-positive cells were excluded due to individual cell quality control performed in multiome analysis (Fig. 3A-B and Supplementary Tables S2 and S3). Of the 27 remaining cells, 8 were detected in ATAC data, 18 in GEX, and 1 in both datasets (Fig. 4B). Infected cells were annotated as 18 CD4^+^ T cells, 6 CD8^+^ T cells, 2 B cells, and 1 monocyte. Both modalities identified CD4^+^ T cells, but GEX uniquely identified HIV-positive B cells, CD8^+^ T cells and the monocyte (Fig. 4C). Importantly, both DNA and RNA were detected in a single CD4^+^ T cell, the primary target of HIV-1. Most HIV-positive cells were still found in post-HIV samples (24 post-HIV versus 3 post-ART); B and CD8^+^ T cells were only recovered in post-HIV samples, monocytes in one individual after one year of ART, and CD4^+^ T cells were observed in both post-infection timepoints and in all participants. Albeit the reduction in the cell numbers, we consistently detected 1-3 viral DNA or RNA counts per cell and still identified the CD4^+^ T cell from HIV8 sample with 119 RNA counts (Fig. 3D). However, the ability of the method to predict proviral levels was diminished (Supplementary Fig. S1C-D).

**Figure 3.**
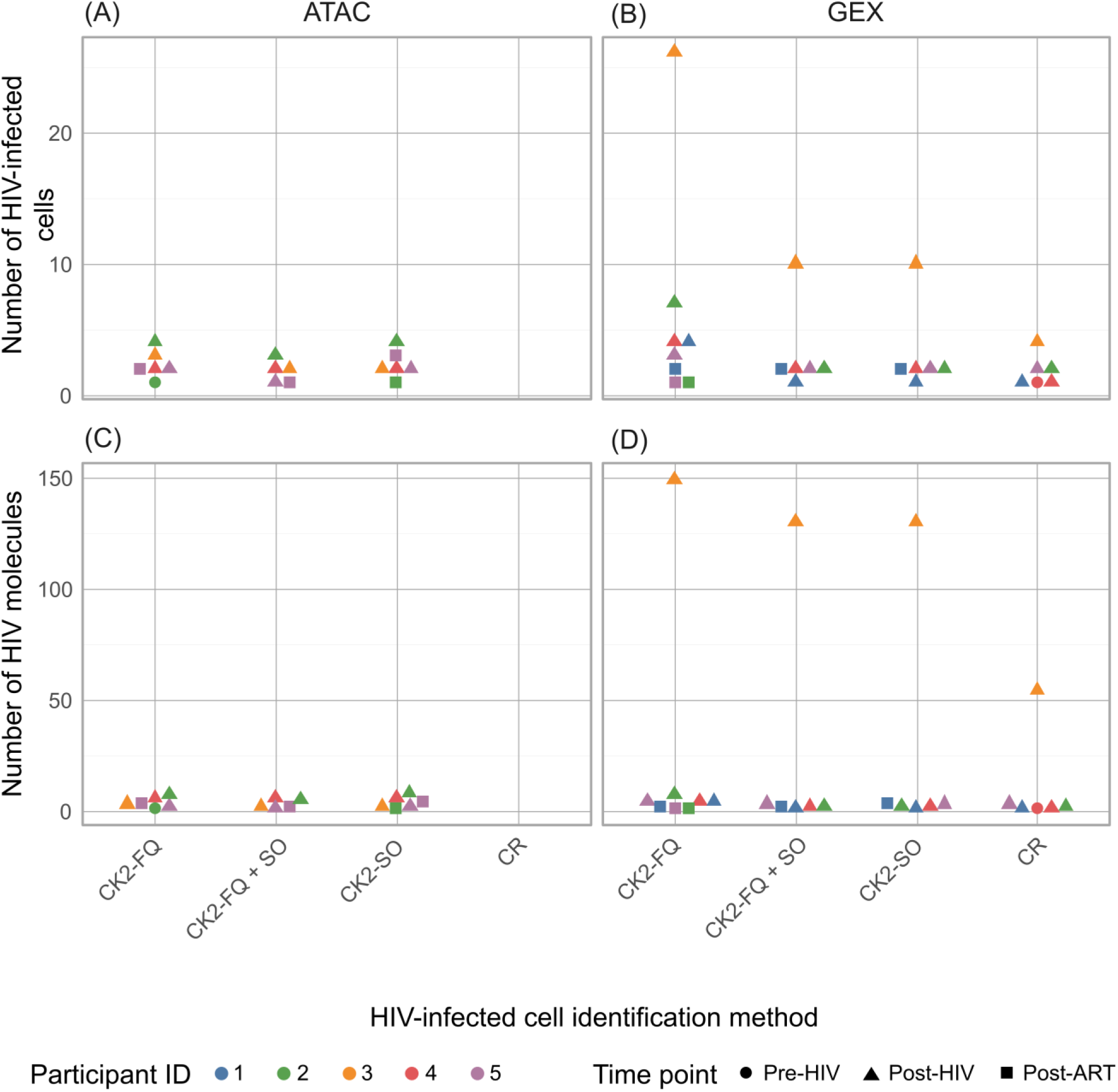
Summary of viral cell and molecule identification reported by K2 and CR. The top row displays the number of HIV-1 infected cells detected using different strategies in ATAC (A) and GEX (B) datasets. The bottom row shows the number of viral DNA molecules (C) and RNA molecules (D) detected.

**Figure 4.**
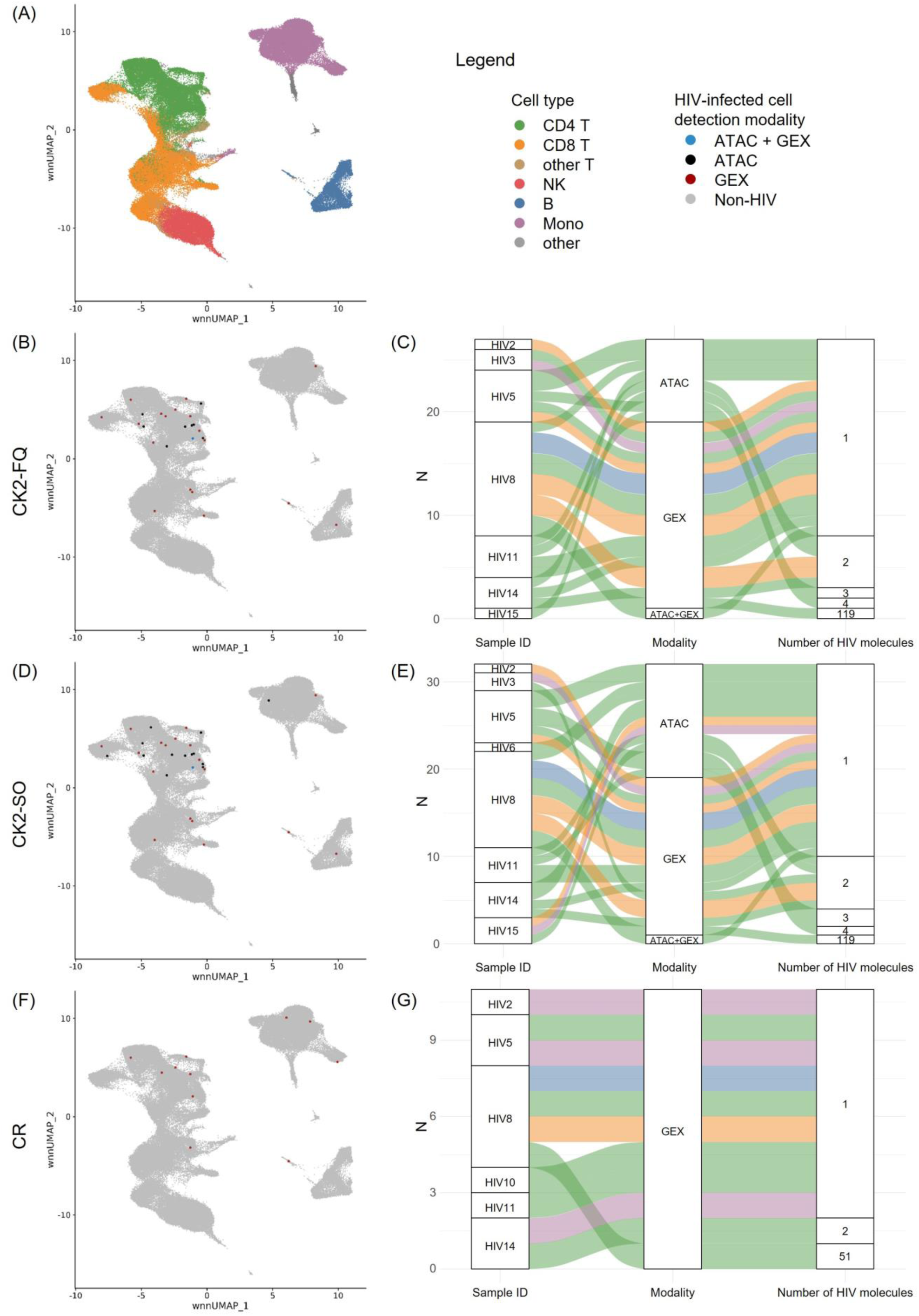
Summary of results for the detection of HIV-1 infected cells and cell types using K2 and CR. First, we retrieved the original single-cell multiome dataset from HIV-scAR project and presented (A) a UMAP representation of individual cells and their cell type. Colors indicate distinct cell types annotated at Azimuth level 1. Next, HIV-infected cells identified using the CK2-FQ calibration strategy were transferred into the Seurat Object from HIV-scAR project (termed as CK2-FQ+SO) as shown in (B). In this panel, individual cells are colored as follows: non-HIV infected cells are shown in gray; infected cells detected with ATAC-seq data are shown in black; infected cells detected with GEX data are shown in red; and cells containing both viral DNA and RNA are indicated in blue. Detailed information for each cell, including sample ID, the modality identifying viral DNA or/and RNA, and the number of viral molecules (i.e.: DNA or/and RNA counts), is visualized as an alluvial plot (C), where linking ribbons represent the number of cells and colors indicate the cell type as in subplot A. Panel (D) display a UMAP showing infected cells identified by CK2-SO strategy. Panel (E) mirrors (C) but for CK2-SO. Panel (F) shows a UMAP with the HIV-infected cells detected by CR. Panel (G) mirrors (C), but for CR.

We further assessed the performance of K2 by tuning its confidence score examining classifications within the Seurat object (termed as CK2-SO). This approach allowed us to lower the confidence threshold, thereby increasing the number of HIV-positive cells and viral reads detected (Fig. 3 and Supplementary Table S3), while maintaining concordance with proviral DNA levels (Supplementary Fig. S1E-F). The ATAC datasets benefited most from this strategy, with the identification of five additional HIV-positive cells containing either one or three DNA counts per cell (Fig. 3 and Fig. 4D-E). These newly detected infected cells were annotated as three CD4^+^ T cells, one CD8^+^ T cell, and one monocyte. In contrast, the GEX datasets reported only a single additional RNA count in an already identified infected CD4^+^ T cell from sample HIV3. Overall, this calibration approach improves the detection and prediction of HIV-1 infected cells compared to previous results obtained at the CK2-FQ and CK2-FQ+SO strategies.

In summary, K2-based identification of viral reads using the CK2-FQ strategy yielded results consistent with observed proviral DNA, but the CK2-SO strategy further improved detection. Regardless of the analytical approach, most infected cells were identified using RNA data from post-HIV samples and were predominantly annotated as CD4^+^ T cells.

### 4.3. Evaluation of HIV-1 positive cells identified by Cell Ranger-ARC

To cross-validate the K2-based detection of HIV-1-positive cells with an alignment-based method, we executed Cell Ranger-ARC (CR) with the hybrid human-HIV_HXB2_ reference. This strategy failed to detect any viral DNA or RNA in any sample. In contrast, when CR was run using all 1,163 curated HIV-1 subtype B reference genomes, we identified viral RNA in 11 cells from six out of 15 samples but did not detect viral DNA (Fig. 3 and Fig. 4F-G). Importantly, one cell was identified in a pre-HIV sample, indicating that CR produced a false positive (FP) both in terms of the number of cells and the number of transcripts detected. The remaining 10 HIV-positive cells were found in post-HIV samples, averaging 2.0 ± 1.2 infected cells per sample. Comparison with proviral DNA measurements revealed poor concordance between these results (Supplementary Fig. S1G-H). Most HIV-positive cells expressed approximately 1-2 viral transcripts per cell, except for sample HIV8, which contained one cell with 51 RNA counts (Fig. 3D, Fig. 4G, and Supplementary Table S3). Cell annotation identified the HIV-positive cells as six CD4^+^ T cells, one CD8^+^ T cell, three monocytes, and one B cell. Notably, both the false positive cell and the transcriptionally activated cell were classified as CD4^+^ T cells. Overall, in our experimental setting, CR yielded an FP detection and showed poor agreement with proviral DNA quantification.

### 4.4. Bioinformatic tools comparison

We next compared the performance of the K2- and CR-based detection approaches. Using default parameters, both bioinformatic tools identified FPs in pre-HIV samples; however, adjusting the confidence scores in K2 effectively eliminated them. Analysis of infected samples revealed that K2 detected a greater number of HIV-positive cells (32 versus 11; Fig. 5A). Seven of these cells were identified by both tools, comprising five CD4^+^ T cells, one CD8^+^ T cell, and one B cell (Supplementary Table S3). Importantly, the transcriptionally active CD4^+^ T cell was detected by both tools, but K2 attributed a higher number of transcripts to this cell (119 transcripts identified by K2 versus 51 by CR). Overall, K2 yielded higher counts of viral DNA and RNA (Fig. 3C-D), and improved concordance with proviral results (Supplementary Figure S1). Despite these differences, Bland-Altman analysis showed concordance between K2 and CR, even after excluding the outlier from sample HIV8 (Figures 5B and 5C). In summary, K2 outperformed CR both in detecting HIV-1-infected cells and in quantifying viral DNA and RNA.

**Figure 5.**
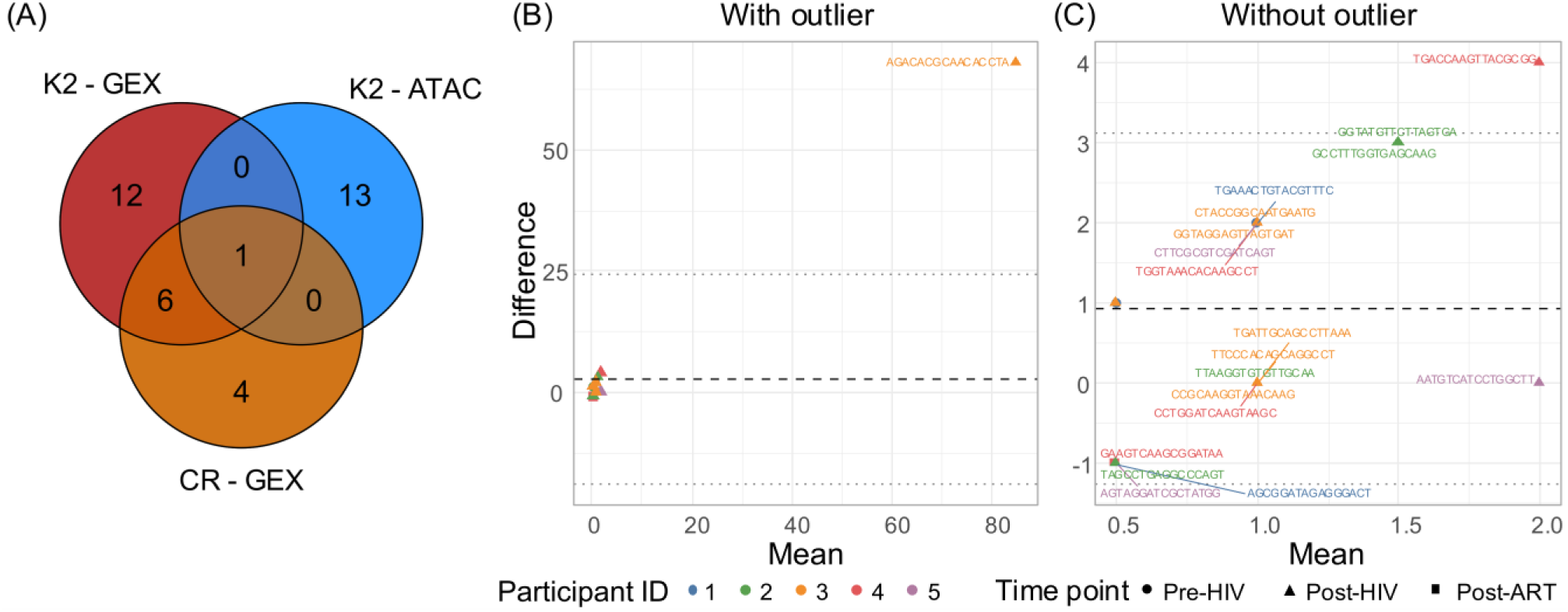
Comparison of results between K2 and CR. (A) Number of infected cells identified by each classifier and modality. (B) Dot plot of Bland-Altman analysis illustrating the number of DNA/RNA molecules detected per cell. Colors indicate the Individual ID, the shape denotes the time point, and the DNA sequences show the cell barcodes. (C) As in panel B but excluding outliers. In panels B and C, dashed lines represent the mean difference, while dotted lines indicate the limits of agreements (calculated as the mean difference ± 1.96 x standard deviation).

Regarding computational efficiency, calibrating K2 required approximately one day (23.3h) when executed with 16 threads on five pre-HIV samples, following the author’s recommendations (27). This process included runs with eleven different confidence score thresholds. Subsequently, K2 was applied to ten samples (post-HIV and post-ART) using two sets of optimized confidence scores (ATAC = 0.30 and GEX = 0.10, and ATAC = 0.15 and GEX = 0), which took around 10h to complete. For each association of viral reads classified by K2 to individual cells, we used our in-house R package, virKra (v1.0.0). Running virKra on a single thread required approximately 6.5h to process both sets of calibrated confidence scores. In total, the integration of K2 into the analysis pipeline required about 40h.

## 5. Discussion

Despite significant advances in single-cell methodologies, reliably identifying HIV-1 infected cells from single-cell data using bioinformatics approaches remains highly challenging. In the context of HIV-research default tools such as CR may be inadequate for detecting infected cells, underscoring the need for alternative computational strategies. In this study, we developed a pipeline focused on the accurate classification of HIV-1 reads, prioritizing detection over genomic location.

In this study, we began by assessing the effect of reference database curation on the identification of viral genomes. When K2 was applied to pre-HIV samples using a non-curated reference database, we detected six viral genomes containing spurious sequences ranging from 30 to 387 nucleotides in length. All of these sequences were located outside the annotated region at either the 5’ or 3’ ends of the reference, indicating that they resulted from inadequate trimming of genome termini or the presence of adaptor sequences. Although elucidating the exact motifs responsible for these problematic sequences falls outside the scope of this study, we excluded them from further analysis. Notably, conventional aligners may miss these short, misassembled fragments when filtering by mapping quality (MAPQ), whereas K2 successfully detected them due to its use of short minimizers (l-mers) of length 31 for k-mers classification. These findings highlight the necessity of curating all genomes prior to analysis to remove misassembled sequences, and demonstrate that K2 is highly sensitive to contamination, making it well-suited for detecting poorly assembled references containing spurious genomic fragments of various lengths.

After thorough curation of the reference database, calibration of K2 was performed independently for each multiome library type (ATAC and GEX) and for both analytical strategies (CK2-FQ versus CK2-SO). For the CK2-FQ calibration, we established the confidence score threshold while excluding the read pair from the HIV4 sample, as these sequences were unequivocally viral in origin. Notably, this read pair shared the same barcode as another infected cell from the post-HIV sample HIV5, both belonging to the same individual and sequenced together in the same flow cell and lane (Supplementary Table S2). This detection was attributed to tag-jumping (index-hopping) and was manually removed (20, 47). A common approach to mitigate such issues is to define a minimum detection threshold of at least 2-5 reads assigned to the virus or unique molecular identifiers (UMIs) (16, 19). However, this recommendation is primarily intended for alignment-based tools operating with default settings, which does not apply to our methodology. In fact, we did not employ the recommended threshold because it exceeds the expected number of HIV-1 reads. Optimizing tools parameters enhances the accuracy of read-to-reference assignments and reduces reliance on stringent threshold-based filtering to eliminate false positives. Ideally, researchers who expect a minimum of two or more UMI (or reads) per sample in the experimental design should apply both strategies (calibration and detection limit) to maximize accurate identifications of HIV-1 reads. However, when viral information is limited, it may be preferable to retain all counts to ensure comprehensive detection.

In our CK2-SO strategy, we reduced confidence scores compared to those used in the previous CK2-FQ approach. This reduction was necessary because some viral reads were lost during the transfer into the Seurat Object using virKra. The underlying causes for this loss include sequencing errors in cell barcodes and dataset filtering. Sequencing errors can alter the sequences used for cell identification, resulting in invalid identifiers. Our virKra package discards all such sequences, whereas CR employs an algorithm based on Hamming distance to correct potential barcode errors (28, 29). However, this issue was observed only with a single pair of reads that exceeded the 0.15 threshold (Supplementary Table S2). Additionally, the loss of infected cells may result from cell filtering steps. The Seurat Object used in this study underwent rigorous pre-processing to remove low-quality cells, doublets, and empty droplets, which may inadvertently exclude actively infected cells. Regardless of the pipeline, our findings and previous studies highlight the necessity of moving beyond default parameters to achieve accurate results in single-cell multiome analyses (16, 40, 43).

Throughout the post-HIV and post-ART phases, we demonstrated that our pipeline reliably reflects proviral status through both viral DNA and RNA detection. Annotation of infected cells further validates our results by enabling cross-verification of whether the identified cells are biologically plausible. Using the Azimuth reference (32), we confirmed that CD4^+^ T cells were the most abundant infected cell type, while we also detected CD8^+^ T cells, B cells, and monocytes. Although CD4^+^ T cells are the primary targets of HIV-1, previous studies have reported the eventual presence of HIV-1 in CD8^+^ T cells and B cells during acute infection (48, 49), and monocytes are increasingly recognized as an underestimated HIV-1 reservoir (48–50). It is important to interpret cell annotation with caution, as misclassification can occur; for example, the CD8^+^ T cell identified in the post-ART HIV15 sample had a prediction score of only 0.7, suggesting possible annotation uncertainty.

This pipeline may also facilitate the distinction between latently infected cells (viral DNA+), productively infected cells (viral RNA+ and optionally viral DNA+), and abortive infections. However, given the low number of HIV-1-infected cells identified and the limited detection of viral DNA by ATAC in our multiome dataset, only limited conclusions can be drawn regarding the infection state or proviral genome intactness. Similar limitations have been reported by Wei and colleagues on their study targeting HIV-1-infected cells on people living with HIV-1 using DOGMA-seq (51).

After aligning the data with CR using two different reference strategies (a single HIV_HXB2_ genome and a comprehensive HIV-1 subtype B database containing 1,163 distinct sequences), we observed that including a broader array of genomes better characterizes HIV-positive cells. Nonetheless, this likely underestimates the true number of infected cells, as CR’s ability to classify reads diminishes with increased multi-mapping; reads aligned to multiple viral genomes are excluded from analysis (http://support.10xgenomics.com/). Similarly, León-Rivera and colleagues (50) analyzed GEX data with CR and evaluated its performance using different HIV-1 genome annotation strategies: annotating the entire genome as a single transcript versus annotating individual genes separately. They report that, although suboptimal, gene-level annotation improved detection sensitivity. In our study, we also tested both approaches, but our preliminary results differed from those of León-Rivera et al.; our identification of viral reads improved when whole genomes were annotated as a single transcript (data not shown). Specifically, when using individual gene annotations, we detected five infected cells, two from pre-HIV samples, each with one viral transcript, and three from the HIV8 sample, with 37, 1 and 1 viral transcripts, respectively. Furthermore, Peterson and colleagues recently used CR with the HIV_HXB2_ genome to identify viral transcripts from people with HIV receiving ART (52). They detected 57 infected CD4^+^ T cells out of 232,936 PBMCs, which contrast with our findings, as we did not detect any HIV DNA+ cell in ART-treated samples with CR. However, these results are not directly comparable, since their samples underwent magnetic CD4^+^ T cell positive selection to increase CD4^+^ T cell frequency. Collectively, these findings underscore that the number of detected cells is influenced by the experimental design and highlight that using CR for HIV-1 infected cell detection requires balancing the need to maximize coverage of viral genetic diversity with maintaining the pipeline’s capacity to accurately classify reads.

The reported results demonstrate the superior performance of K2 compared to CR when detecting HIV-1 subtype B molecules (both DNA and RNA) in single-cell datasets. Notably, K2 was optimized through rigorous database curation and confidence score tuning, whereas CR was not calibrated for these reasons (refer to the Materials and Methods section). Integrating this pipeline into the analysis workflow increases computational demands, as users must tailor K2 settings using negative or positive controls. Therefore, researchers should evaluate whether additional analysis will benefit their study objectives. Our virKra package, developed to facilitate the transfer of K2 outputs into Seurat objects, is designed for user accessibility, although current version introduces analytical complexity and does not include a graphical user interface. Data storage requirements are minimal: the K2 reference database and output files together require only 34.3 MB (3.3 MB for the reference database, 16 MB for ATAC output, and 15 MB for GEX).

## 6. Conclusions

In summary, this study introduces a novel bioinformatic pipeline for the identification of HIV-positive cells, employing a calibrated Kraken2 (K2) classifier integrated with a set of multiple subtype B reference genomes. This strategy enhances the detection of HIV-infected cells compared to Cell Ranger-ARC (CR) across both single-cell multiome modalities (ATAC and GEX). Notably, while CR generated a false positive in a pre-HIV sample, our K2-based pipeline detected infected cells exclusively in HIV-positive samples. This pipeline holds promise as a valuable tool for studies seeking to identify cellular markers specific to HIV-infected cells.

## 7. Declarations

### Ethics approval and consent to participate

Participants gave written informed consent to participate in accordance with the Declaration of Helsinki. The study was approved by the local independent institutional review boards at Barcelona’s Clinic hospital and Germans Trias i Pujol Hospital (Check-Ear: EO-09-014, and CHAVI: EO-08-023).

## Consent for publication

JM-P as co-corresponding author confirms that all authors have read and agreed with the current version of the manuscript.

## Availability of data and materials

ATAC-Seq data was deposited to European Genome-phenome Archive (EGA) with accession number available upon acceptance.

RNA-Seq data was deposited to European Genome-phenome Archive (EGA) with accession number available upon acceptance.

## Authors’ contributions

LG-S, JM-P, and SM-L conceived the idea. LG-S, LBS and SM-L performed the data curation. LG-S and SM-L designed the methodology; LG-S performed the analysis and VU helped with the statistics analysis. CB, PC and BM contributed to the cohort design and management and clinical data collection. LG-S, SM-L, and JM-P were responsible for the decision to submit the manuscript. SM-L, VU, RL-R, LBS, MCP, JD, PC, BM, CB, BC, AT and JM-P made important critical reviews and feedback on the manuscript. JM-P, as co-corresponding author, confirms that all authors have read and agreed with the current version of the manuscript. The co-corresponding author had full access to all the data in the study and had final responsibility for the decision to submit for publication.

## Acknowledgments

Firstly, we thank all the participants and BCN Checkpoint, without whom this study could not have been performed. We are grateful to the technical staff of IrsiCaixa for sample processing (L. Ruiz, E. Grau, R. Ayen, L. Gomez, C. Ramirez, M. Martinez, and T. Puig).

## 9. Supporting information captions

Figure S1. Comparison between expected and estimated viral DNA/RNA quantities. The graphs on the left (A, C, E, and G) display the data distribution, with the black line indicating the 1:1 ratio. The graphs on the right (B, D, F, and H) present Bland-Altman analyses, both including and excluding the outlier sample (i.e. HIV8). Panels A-B show results for calibrated Kraken2 (CK2-FQ method) using a confidence score of 0.30 for ATAC and 0.10 for GEX. Panels C-D depict results after importing A-B data into the Seurat object (CK2-FQ + SO method). Panels E-F illustrate results with Kraken2 confidence scores adjusted at CK2-SO using confidence scores of 0.15 for ATAC and 0 for GEX. Panels G-H show results from Cell Ranger-ARC (CR method).

Table S1. List of HIV-1 subtype B genomes used in the study. All HIV-1 subtype B genomes downloaded from the LANL HIV database are listed, including their sequence names, accession numbers, whether they were removed during data curation, and, if applicable, the reason for removal.

Table S2. Summary of ATAC and GEX reads classified as HIV-1 by Kraken2. For each read or read pair, we provide the library type (ATAC or GEX), sample, read name, barcodes, barcode quality, UMI sequences, read length, read sequence and quality, Kraken2 output information, and related Seurat object data.

Table S3. Summary of viral molecules (DNA/RNA) detected across different strategies. For each cell, we indicate the sample, barcode, modality (ATAC vs GEX), predicted cell type, and the number of molecules detected, separated by detection strategy.

Table S4. Laboratory and bioinformatic results of viral RNA and DNA data.

## References

1. Maartens, G., Celum, C. and Lewin, S.R. (2014) HIV infection: epidemiology, pathogenesis, treatment, and prevention. The Lancet, 384, 258–271.

2. Preston, B.D., Poiesz, B.J. and Loeb, L.A. (1988) Fidelity of HIV-1 reverse transcriptase. Science, 242, 1168–1171.

3. Roberts, J.D., Bebenek, K. and Kunkel, T.A. (1988) The accuracy of reverse transcriptase from HIV-1. Science, 242, 1171–1173.

4. Mourez, T., Simon, F. and Plantier, J.-C. (2013) Non-M variants of human immunodeficiency virus type 1. Clin Microbiol Rev, 26, 448–461.

5. Indihar, D.F., Oke, R., Hemerding, R., et al. (2025) Rare but relevant: a comprehensive narrative review of rare HIV-1 group M subtypes. mBio, 16, e01264–25.

6. Abecasis, A.B., Wensing, A.M., Paraskevis, D., et al. (2013) HIV-1 subtype distribution and its demographic determinants in newly diagnosed patients in Europe suggest highly compartmentalized epidemics. Retrovirology, 10, 7.

7. Robertson, D.L., Anderson, J.P., Bradac, J.A., et al. (2000) HIV-1 nomenclature proposal. Science, 288, 55–55.

8. Iyer, L.R. and Thomas, R. (2025) Current insight into HIV-1 persistence from single-cell transcriptome profiling in acutely treated cohorts of infection. Current Opinion in HIV and AIDS, 20, 481–487.

9. Mbhele, N., Chimukangara, B., Tyers, L., et al. (2025) Advancements in single-cell techniques for examining the HIV reservoir: pathways to a cure. mBio, 16, e00655–25.

10. Peluso, M.J., Bacchetti, P., Ritter, K.D., et al. (2020) Differential decay of intact and defective proviral DNA in HIV-1–infected individuals on suppressive antiretroviral therapy. JCI Insight, 5, e132997.

11. Simonetti, F.R., White, J.A., Tumiotto, C., et al. (2020) Intact proviral DNA assay analysis of large cohorts of people with HIV provides a benchmark for the frequency and composition of persistent proviral DNA. Proc. Natl. Acad. Sci. U.S.A., 117, 18692–18700.

12. Chun, T.-W., Stuyver, L., Mizell, S.B., et al. (1997) Presence of an inducible HIV-1 latent reservoir during highly active antiretroviral therapy. Proc. Natl. Acad. Sci. U.S.A., 94, 13193–13197.

13. Finzi, D., Hermankova, M., Pierson, T., et al. (1997) Identification of a reservoir for HIV-1 in patients on highly active antiretroviral therapy. Science, 278, 1295–1300.

14. Wong, J.K., Hezareh, M., Günthard, H.F., et al. (1997) Recovery of Replication-Competent HIV Despite Prolonged Suppression of Plasma Viremia. Science, 278, 1291–1295.

15. Sierra, S., Kupfer, B. and Kaiser, R. (2005) Basics of the virology of HIV-1 and its replication. Journal of Clinical Virology, 34, 233–244.

16. Wong, M., Wei, Y. and Ho, Y.-C. (2023) Single-cell multiomic understanding of HIV-1 reservoir at epigenetic, transcriptional, and protein levels. Current Opinion in HIV and AIDS, 18, 246– 256.

17. Li, H. and Durbin, R. (2009) Fast and accurate short read alignment with Burrows–Wheeler transform. Bioinformatics, 25, 1754–1760.

18. Dobin, A., Davis, C.A., Schlesinger, F., et al. (2013) STAR: ultrafast universal RNA-seq aligner. Bioinformatics, 29, 15–21.

19. Collora, J.A., Liu, R., Pinto-Santini, D., et al. (2022) Single-cell multiomics reveals persistence of HIV-1 in expanded cytotoxic T cell clones. Immunity, 55, 1013–1031.e7.

20. De Vries, J.J.C., Brown, J.R., Couto, N., et al. (2021) Recommendations for the introduction of metagenomic next-generation sequencing in clinical virology, part II: bioinformatic analysis and reporting. Journal of Clinical Virology, 138, 104812.

21. Lee, D., Choi, S.Y., Shin, S.-I., et al. (2025) Multi-omics single-cell analysis reveals key regulators of HIV-1 persistence and aberrant host immune responses in early infection. eLife, 14, RP104856.

22. Bost, P., Giladi, A., Liu, Y., et al. (2020) Host-viral infection maps reveal signatures of severe COVID-19 patients. Cell, 181, 1475–1488.e12.

23. Zhang, W., Xu, X., Fu, Z., et al. (2022) PathogenTrack and Yeskit: tools for identifying intracellular pathogens from single-cell RNA-sequencing datasets as illustrated by application to COVID-19. Front. Med., 16, 251–262.

24. Lee, C.Y., Chen, Y., Duan, Z., et al. (2022) Venus: An efficient virus infection detection and fusion site discovery method using single-cell and bulk RNA-seq data. PLoS Comput Biol, 18, e1010636.

25. Whitmore, L.S., Tisoncik-Go, J. and Gale, M. (2024) scPathoQuant: a tool for efficient alignment and quantification of pathogen sequence reads from 10× single cell sequencing datasets. Bioinformatics, 40, btae145.

26. Wu, L.-Y., Wijesekara, Y., Piedade, G.J., et al. (2024) Benchmarking bioinformatic virus identification tools using real-world metagenomic data across biomes. Genome Biol, 25, 97.

27. Wood, D.E., Lu, J. and Langmead, B. (2019) Improved metagenomic analysis with Kraken 2. Genome Biol, 20, 257.

28. Zheng, G.X.Y., Terry, J.M., Belgrader, P., et al. (2017) Massively parallel digital transcriptional profiling of single cells. Nat Commun, 8, 14049.

29. Satpathy, A.T., Granja, J.M., Yost, K.E., et al. (2019) Massively parallel single-cell chromatin landscapes of human immune cell development and intratumoral T cell exhaustion. Nat Biotechnol, 37, 925–936.

30. Moron-Lopez, S., Garrido-Sanz, L., Soriaga, L., et al. (2026) Longitudinal multiomics analysis in blood from individuals before and after HIV acquisition. bioRxiv, 10.64898/2026.11.23.767218.

31. Andrews, S. (2015) FastQC: a quality control tool for high throughput sequence data. URL: http://www.bioinformatics.babraham.ac.uk/projects/fastqc

32. Hao, Y., Hao, S., Andersen-Nissen, E., et al. (2021) Integrated analysis of multimodal single-cell data. Cell, 184, 3573–3587.e29.

33. Stuart, T., Srivastava, A., Madad, S., et al. (2021) Single-cell chromatin state analysis with Signac. Nat Methods, 18, 1333–1341.

34. R Core Team (2023) R: A language and environment for statistical computing. URL: https://www.R-project.org/

35. Young, M.D. and Behjati, S. (2020) SoupX removes ambient RNA contamination from droplet-based single-cell RNA sequencing data. GigaScience, 9, giaa151.

36. Germain, P.-L., Lun, A., Garcia Meixide, C., et al. (2022) Doublet identification in single-cell sequencing data using scDblFinder. F1000Res, 10, 979.

37. Zhang, Y., Liu, T., Meyer, C.A., et al. (2008) Model-based analysis of ChIP-seq (MACS). Genome Biol, 9, R137.

38. Stuart, T., Butler, A., Hoffman, P., et al. (2019) Comprehensive integration of single-cell data. Cell, 177, 1888–1902.e21.

39. Pilo, P., Lawless, C., Tiley, A.M.M., et al. (2022) Comparison of microscopic and metagenomic approaches to identify cereal pathogens and track fungal spore release in the field. Front. Plant Sci., 13, 1039090.

40. Garrido-Sanz, L., Senar, M.À. and Piñol, J. (2022) Drastic reduction of false positive species in samples of insects by intersecting the default output of two popular metagenomic classifiers. PLoS ONE, 17, e0275790.

41. Peabody, M.A., Van Rossum, T., Lo, R., et al. (2015) Evaluation of shotgun metagenomics sequence classification methods using in silico and in vitro simulated communities. BMC Bioinformatics, 16, 362.

42. Lupo, V., Van Vlierberghe, M., Vanderschuren, H., et al. (2021) Contamination in reference sequence databases: time for divide-and-rule tactics. Front Microbiol, 12, 755101.

43. Liu, R., Yeh, Y.-H.J., Varabyou, A., et al. (2020) Single-cell transcriptional landscapes reveal HIV-1–driven aberrant host gene transcription as a potential therapeutic target. Sci. Transl. Med., 12, eaaz0802.

44. Altschul, S.F., Gish, W., Miller, W., et al. (1990) Basic local alignment search tool. Journal of Molecular Biology, 215, 403–410.

45. Zhang, Z., Schwartz, S., Wagner, L., et al. (2000) A greedy algorithm for aligning DNA sequences. Journal of Computational Biology, 7, 203–214.

46. Shen, W., Le, S., Li, Y., et al. (2016) SeqKit: a cross-platform and ultrafast toolkit for FASTA/Q file manipulation. PLoS ONE, 11, e0163962.

47. Sinha, R., Stanley, G., Gulati, G.S., et al. (2017) Index switching causes “spreading-of-signal” among multiplexed samples in Illumina HiSeq 4000 DNA sequencing. bioRxiv, 10.1101/125724.

48. Wu, V.H., Nordin, J.M.L., Nguyen, S., et al. (2023) Profound phenotypic and epigenetic heterogeneity of the HIV-1-infected CD4+ T cell reservoir. Nat Immunol, 24, 359–370.

49. Delley, C.L., Shah, S., Joslin, K.M., et al. (2025) The immunophenotype and proviral landscape of HIV-infected CD4 T cells during antiretroviral therapy. bioRxiv, 10.1101/2025.06.05.657210.

50. León-Rivera, R., Morsey, B., Niu, M., et al. (2020) Interactions of monocytes, HIV, and ART identified by an innovative scRNAseq pipeline: pathways to reservoirs and HIV-associated comorbidities. mBio, 11, e01037–20.

51. Wei, Y., Davenport, T.C., Collora, J.A., et al. (2023) Single-cell epigenetic, transcriptional, and protein profiling of latent and active HIV-1 reservoir revealed that IKZF3 promotes HIV-1 persistence. Immunity, 56, 2584–2601.e7.

52. Peterson, J.J., Chandel, S., James, K., et al. (2025) Single cell characterization of the gastrointestinal HIV reservoir reveals heterogeneous cellular phenotypes. Journal of Clinical Investigation, 10.1172/JCI196536.

