## Supplemental Figure 1 for "Identification of HIV-1 infected cells in single-cell multiome datasets using Kraken2"

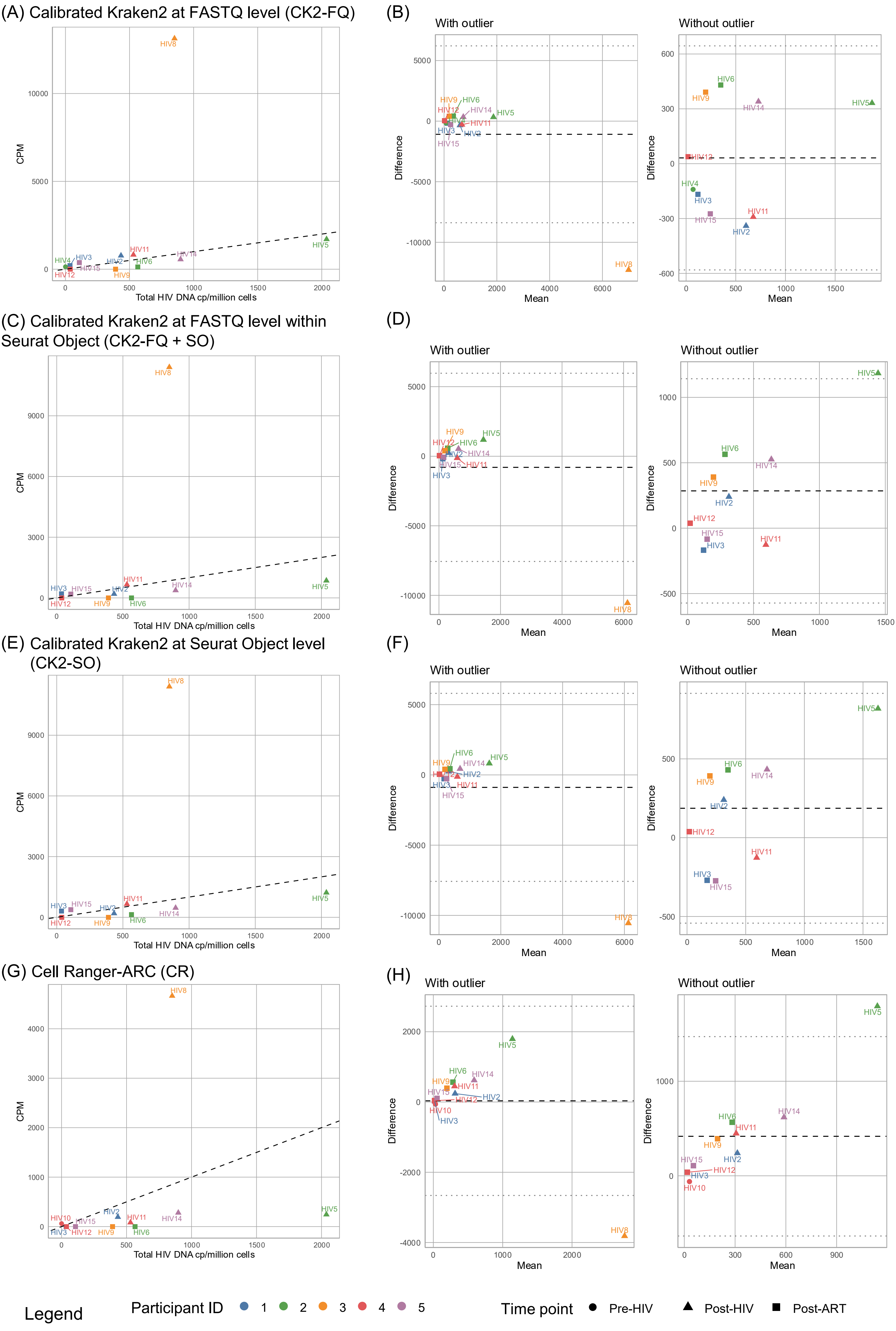
(A) Calibrated Kraken2 at FASTQ level (CK2-FQ)
CPM
Total HIV DNA cp/million cells
(B)
With outlier
Without outlier
Difference
Mean
(C) Calibrated Kraken2 at FASTQ level within Seurat Object (CK2-FQ + SO)
CPM
Total HIV DNA cp/million cells
(D)
With outlier
Without outlier
Difference
Mean
(E) Calibrated Kraken2 at Seurat Object level (CK2-SO)
CPM
Total HIV DNA cp/million cells
(F)
With outlier
Without outlier
Difference
Mean
(G) Cell Ranger-ARC (CR)
CPM
Total HIV DNA cp/million cells
(H)
With outlier
Without outlier
Difference
Mean
HIV2
HIV3
HIV4
HIV5
HIV6
HIV8
HIV9
HIV10
HIV11
HIV12
HIV14
HIV15
Legend
Participant ID 1 2 3 4 5
Time point Pre-HIV Post-HIV Post-ART

**Supplementary Figure 1.** Comparison between expected and estimated viral DNA/RNA quantities. The graphs on the left (A, C, E, and G) display the data distribution, with the black line indicating the 1:1 ratio. The graphs on the right (B, D, F, and H) present Bland-Altman analyses, both including and excluding the outlier sample (i.e. HIV8). Panels A-B show results for calibrated Kraken2 (CK2-FQ method) using a confidence score of 0.30 for ATAC and 0.10 for GEX. Panels C-D depict results after importing A-B data into the Seurat object (CK2-FQ + SO method). Panels E-F illustrate results with Kraken2 confidence scores adjusted at CK2-SO using confidence scores of 0.15 for ATAC and 0 for GEX. Panels G-H show results from Cell Ranger-ARC (CR method).
